# *In vivo* adenine base editing corrects newborn murine model of Hurler syndrome

**DOI:** 10.1101/2021.10.16.464213

**Authors:** Jing Su, Xiu Jin, Kaiqin She, Yi Liu, Xiaomei Zhong, Qinyu Zhao, Jianlu Xiao, Ruiting Li, Hongxin Deng, Yang Yang

**Author notes:** Corresponding authors: Yang Yang, State Key Laboratory of Biotherapy and Cancer Center, West China Hospital, Sichuan University and Collaborative Innovation Center, Chengdu 610041, China, Postal address: Ke-yuan Road 4, No. 1, Gao-peng Street, Chengdu, Sichuan, 610041, China. These authors contributed equally: Jing Su, Xiu Jin, Kaiqin She.

## Abstract

Mucopolysaccharidosis type I (MPS I) is a severe disease caused by loss-of-function mutations variants in the α-L-iduronidase (*IDUA*) gene. In *vivo* genome editing represents a promising strategy to correct *IDUA* mutations, and has the potential to permanently restore IDUA function over the lifespan of the patients. Here, we used adenine base editing to directly convert A>G (TAG>TGG) in newborn murine model harboring *Idua-W392X* mutation, which recapitulates the human condition and is analogous to the highly prevalent human W402X mutation. We engineered a split-intein dual-adeno-associated virus (AAV) 9 *in vivo* adenine base editor to circumvent the package size limit of AAV vectors. Intravenous injection of AAV9-base editor system into MPS I newborn mice led to sustained enzyme expression sufficient for correction of metabolic disease (GAGs substrate accumulation) and prevention of neurobehavioral deficits. We observed a reversion of the W392X mutation in 22.46±6.74% of hepatocytes, 11.18±5.25% of heart and 0.34±0.12% of brain, along with decreased GAGs storage in peripheral organs (liver, spleen, lung and kidney). Collectively, these data showed the promise of a base editing approach to precisely correct a common genetic cause of MPS I *in vivo* and could be broadly applicable to the treatment of a wide array of monogenic diseases.

## Introduction

Mucopolysaccharidosis type I (MPS I) is a severe metabolic disorder caused by deficiency of the lysosomal enzyme, α-L-iduronidase (IDUA), which can catalyze the degradation of glycosaminoglycans (GAGs) heparan and dermatan sulfates. The accumulation of GAGs leads to multi-systemic pathologies and diverse clinical manifestations, including cardiomyopathy, hepatosplenomegaly, upper airway obstruction and progressive neurological disease (1, 2). According to the severity of the disease, MPS I was classified as mild (Scheie syndrome or MPS IS; MIM#607016), moderate (Hurler–Scheie syndrome or MPS IH/S; MIM#607015) and severe (Hurler syndrome or MPS IH; MIM#607014) subtypes. MPS IH occurs in approximately 1 in 100,000 newborns and is caused by a variation in the *IDUA* gene (3-5). So far, more than 200 pathogenic variants have been reported, including splicing mutations, insertion and deletions, and missense/nonsense mutations (4). One of the most common mutations (G→A; W402X) accounts for over 40% of patients (6-8).

MPS IH patients begin to show signs of disease within the first 6 months after birth, and will usually die within the first decade without treatment (9). Therefore, early diagnosis and treatment are essential to prevent the development of serious manifestations. Current approved treatments include enzyme replacement therapy (ERT) and hematopoietic stem cell transplantation (HSCT) (10). HSCT is considered standard of care for MPS IH patients, but its success depends on early treatment. Although these treatments can significantly improve disease outcomes and prolong life, there is still a considerable disease burden (11). Many MPS I-related gene therapies and gene editing approaches are under investigation. Studies have reported that *IDUA* gene was delivered to large animals (dog, cat, and rhesus macaques) by AAV through systemic administration or intrathecal injection, effectively alleviating liver, cardiovascular and brain disease phenotypes (12-14). In addition, AAV-mediated zinc finger nucleases and proprietary system gene editing also increased the expression of IDUA *in vivo* and decreased the GAGs storage in MPS I mice (*Idua*^-/-^) (15, 16). Although some promising gene therapy and gene editing results have been obtained in animal models, gene therapy may cause potential loss of an episomal transgene and gene editing may cause unwanted deletion-insertion mutagenesis due to DNA double-strand breaks (DSBs). Base editing can directly convert targeted base pairs without generating DSBs and with minimal indels, so it is considered more suitable for the treatment of human monogenetic inheritance (17, 18). The adenine base editors (ABEs) can convert A•T to G•C, which is composed of dCas9 and adenine deaminase. We reasoned that ABEs can effectively correct MPS IH with G>A point mutation.

In this study, we have developed an AAV9-mediated ABE to directly convert A>G (TAG>TGG) in newborn murine model harboring *Idua*-W392X mutation, which recapitulates the human condition and is analogous to the highly prevalent human W402X mutation. We found partial correction of the pathogenic mutation, biochemical and neurobehavioral deficits in MPS I mice 12 weeks after treatment. These findings provide a potential therapeutic approach for MPS I through directly correction of pathogenic mutations *in vivo*, informing the application of base editing strategies in the treatment of monogenetic disorders.

## Results

### *In vitro* screening of *Idua*-W392X ABEs

*Idua*-W392X mouse is a knock-in disease model of MPS IH that introduces a nucleotide change into the mouse *Idua* gene (19), resulting in a nonsense mutation (G>A) in codon W392 (Figure 1A). To evaluate the base editing efficiency *in vitro*, we generated a HEK293-*Idua* mutant cell lines by stably integrating *Idua*-W392X sequence into AAVS1 genomic locus using CRISPR/Cas9 (Supplemental Figure 1 and Supplemental Table 1). We first searched for protospacer sequences that span the targeted base. We identified two protospacer-adjacent motif (PAM) sites that allowed binding of the corresponding adenine base editors, VRQR-ABEmax, xCas9(3.7)-ABE(x7.10), NG-ABEmax, NG-ABE8e and ABEmax(7.10)-SpG (Figure 1B). We also engineered an ABE8e-SpG similar to NG-ABE8e base editors recently described by Richter et al (20). It was previously reported that these ABEs favorably deaminate within the window of protospacer positions 4-6 (21). We co-transfected these adenine base editors with sgRNA-A5 or sgRNA-A6 into HEK293-*Idua* mutant cell lines to screen the most effective base editors 72h after transfection, respectively. Sanger sequencing showed that sgRNA-A6 had higher on-target editing efficiency than sgRNA-A5, and the editing efficiency of VRQR-ABEmax, xCas9(3.7)-ABE(x7.10), NG-ABEmax, ABEmax(7.10)-SpG, NG-ABE8e and ABE8e-SpG were 3.67±1.53%, 25.67± 4.73%, 5.33±2.01%, 17.33±2.52%, 11.00±1.00% and 39.00±11.53%, respectively (Figure 1C). The highest on-target editing efficiency and no bystander editing were observed with ABE8e-SpG co-transfected with sgRNA-A6, which was then selected for further studies (Figure 1C and 1D). Then, we engineered a split-intein dual-AAV system to effectively deliver the base editors to mice *in vivo*, each expressing one half of the base editor (referred to as N-ABE8e.SpG and C-ABE8e.SpG.sgRNA-A6). The *in vitro* results indicated that the editing efficiency of the split-intein dual-AAV system was lower than that of the full-length ABE8e-SpG version which might be due to a lower abundance of the reconstituted base editor (Figure 1E and 1F).

**Figure 1.**
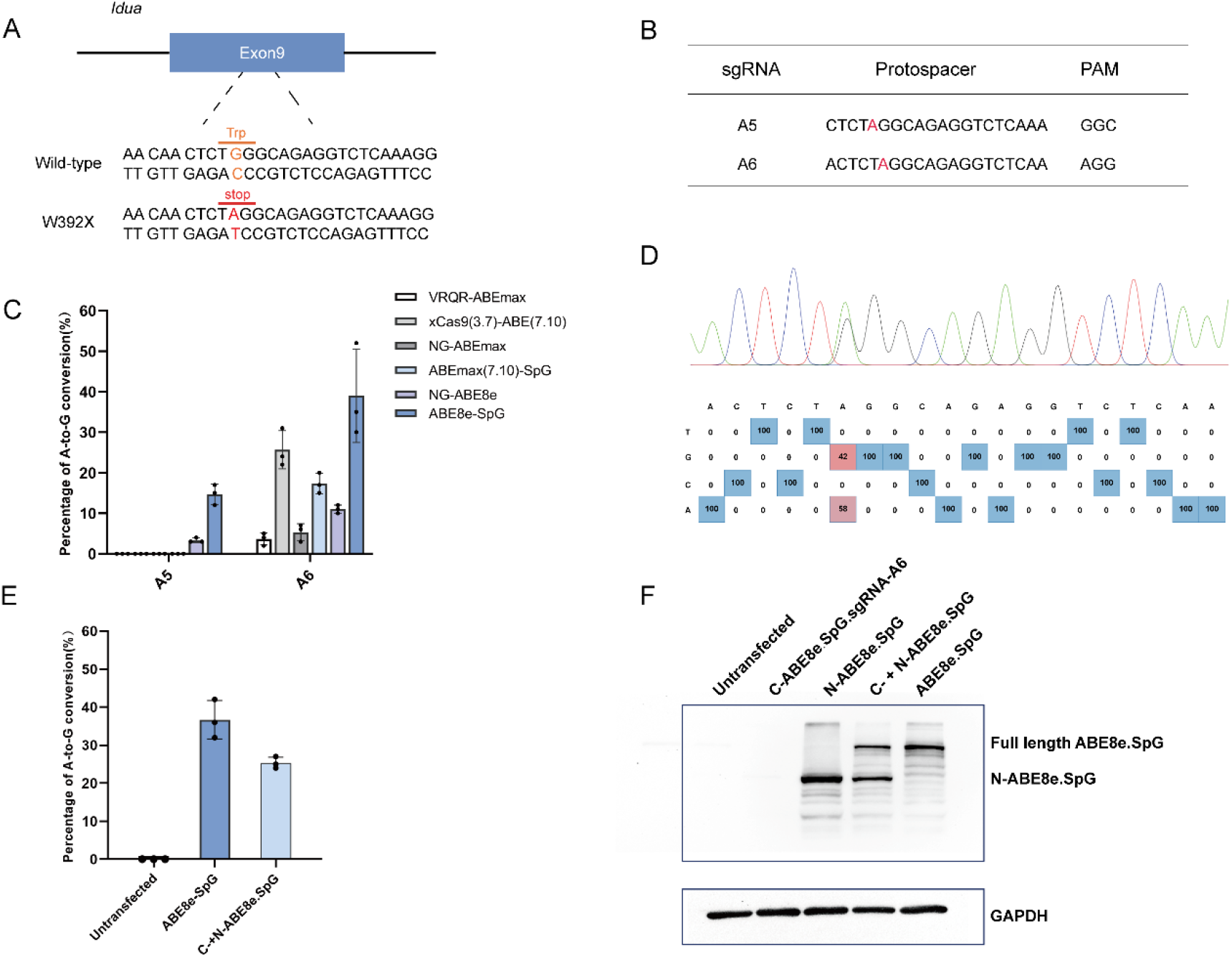
*In vitro* validation of W392X mutation correction by the ABE. (A) The *Idua-*W392X mice has a homozygous G•C to A•T nonsense mutation in exon 9 of the *Idua* gene, changing tryptophan (orange) to a stop codon (red). (B) sgRNA-A5 and sgRNA-A6 were designed to target mutation site (red letter) in the editing window of ABEs. (C) Sanger sequencing analysis of correction efficiency of ABEs in mutant cell lines. (D) Analysis of ABE8e-SpG and sgRNA-A6 co-transfection producing bystander editing. The red grid is the target site. (E) Sanger sequencing analysis of split-intein ABE8e-SpG correction efficiency in mutant cell lines. Transfection of full-length ABE8e-SpG serves as control (n=3 biological replicates each). Mean ± SD are shown. (F) Western blot analysis of co-transfected split-intein ABE8e-SpG. The SpCas9 epitope is only detected at the N-terminal part of the base editor.

### *In vivo* base editing corrects the W392X mutation in the newborn MPS I mice

Since MPS IH is a multi-system disease, we chose the AAV9 serotype for its broad tissue tropism to package the split-intein base editors (refer to as AAV9.N-ABE8e-SpG and AAV9.C-ABE8e-SpG) (Figure 2A) (22, 23). We performed temporal vein injection with AAV9.N-ABE8e-SpG (3×10 ^11^GC/mouse) and AAV9.C-ABE8e-SpG (3×10 ^11^GC/mouse) in newborn *Idua*-W392X mice (n=5). Twelve weeks after AAV treatment, the mice were sacrificed, and the editing efficiency was evaluated in various tissues (Figure 2A). Next-generation sequencing (NGS) results showed effective correction in heart (11.18±5.25%) and liver (22.46±6.74%), and low-level correction in the spleen (0.17±0.02%), lung (0.25± 0.02%), kidney (0.25±0.05%), brain (0.34±0.12%) and muscle (0.30±0.09%) (Figure 2C). Consistent with the NGS results, tissue biodistribution data revealed high AAV9 transduction in the heart (19.89±9.20 GC/cell) and liver (16.35±6.73 GC/cell), with copy numbers below 2.50 GC/cell in other tissues (Figure 2D).

**Figure 2.**
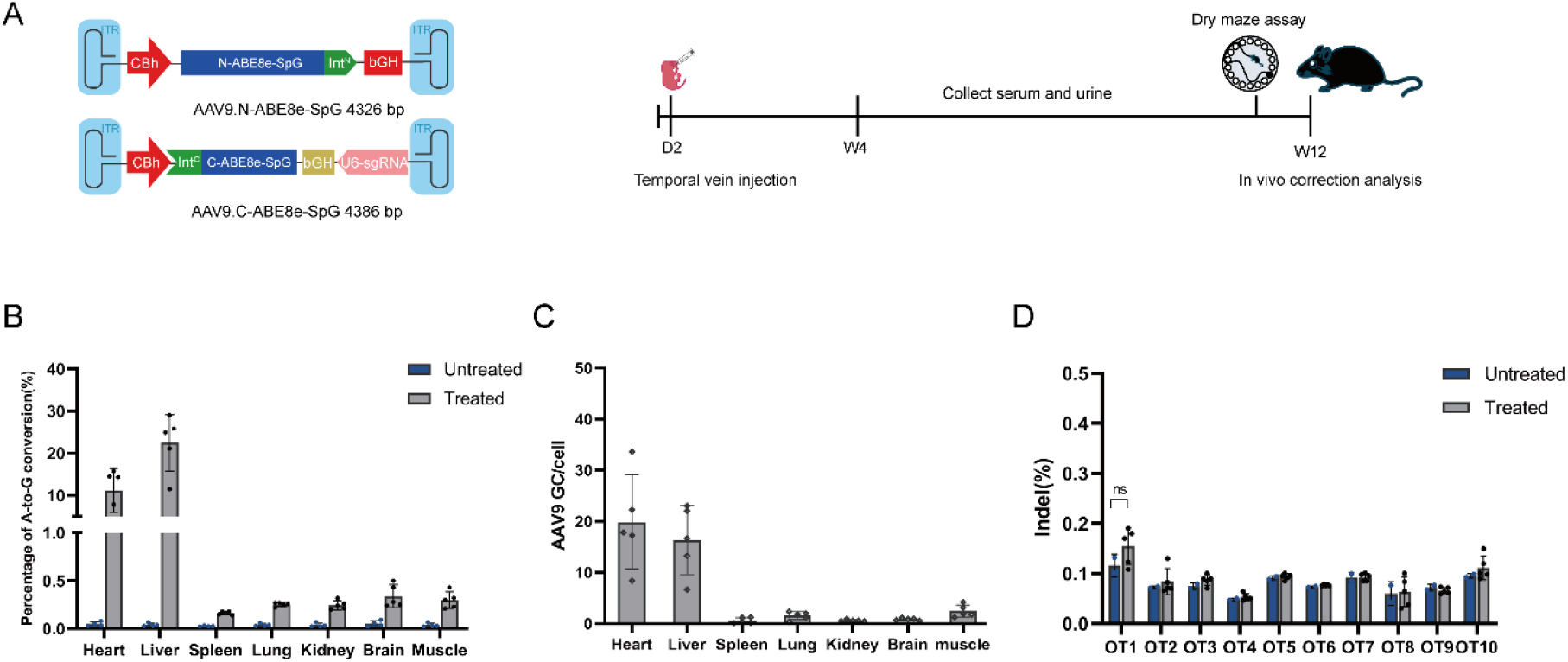
Correction of the pathogenic point mutation in newborn MPS I mice by ABE. (A) Schematic diagram of the genomes of two AAV viral vectors encoding split-intein ABE8e-SpG (left) and a summary of the *in vivo* experiments (right). (B) The correction efficiency of pathogenic mutations in mouse tissue genomic DNA was detected by NGS. Untreated *Idua-*W392X mice (n=4) were included as control. Treated *Idua-*W392X mice (n=5). (C) Quantitative analysis of viral genome copy number in various tissues at 12 weeks post-injection by qPCR. (D) NGS analysis of the top 10 potential off-target sites in liver DNA samples. Untreated *Idua-*W392X mice (n=2) were included as control. Treated *Idua*-W392X mice (n=5). Mean ± SD are shown. Dunnett’s test. ns=non-significant.

Off-target mutations of base editors in genomic DNA may be sgRNA-dependent, caused by the binding of the Cas9-deaminase complex to sequences similar to the target site. Therefore, the algorithm described in www.benchling.com identified the top 10 potential off-target sites for sgRNA-A6 (Supplemental Table 2). These off-target sites were amplified by nest PCR from the liver tissue genomic DNA and deep sequenced with NGS (Supplemental Table 3). We observed similar indel rates in treated mice to untreated mice in these sites and found no evidence of substantial off-target mutations production in genomic DNA after base editing *in vivo* (Figure 2E). Together, these results suggest that adenine base editing is effective and safe *in vivo*.

### *In vivo* base editing increases IDUA enzyme activity and decreases GAGs storage in newborn MPS I mice

In MPS IH patients and *Idua*-W392X mice, there is almost no IDUA enzyme, and thus GAGs accumulate in urine and tissues (24). Serum was collected weekly from 4 weeks after injection to evaluate IDUA enzyme activity. We observed that the serum IDUA enzyme activity of the untreated mice was lower than 0.68 nmol/ml/hr, and the serum IDUA enzyme activity of the treated mice was maintained at about 23.11% of wild-type C57BL/6J mice (mean of 1.77 and 7.66 nmol/ml/hr, respectively) at multiple time points throughout the study period (Figure 3A). The urine GAGs level in the treated mice was about 60% lower than that in the untreated mice (mean of 4.14 and 10.24 mg GAGs/mg Creatinine, respectively) at 12 weeks post injection (Figure 3B).

**Figure 3.**
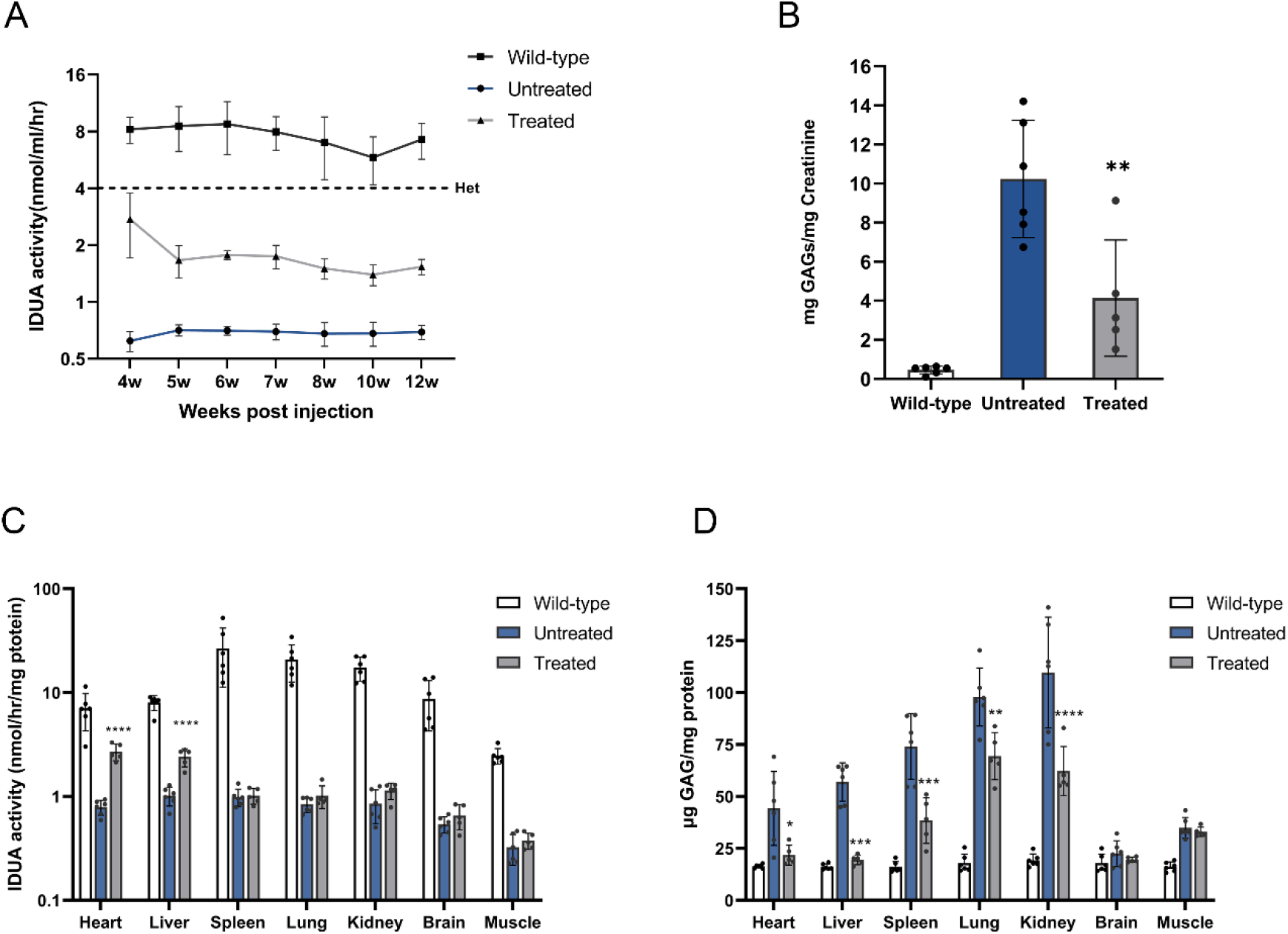
*In vivo* base editing enables sustained biochemical correction in newborn MPS I mice. (A) Time course of serum IDUA activity was measured 4 weeks after injection. Dotted line indicates the serum IDUA activity of heterozygous *Idua*-W392X mice. (B) Urine GAGs was detected 12 weeks after injection. (C) Tissue IDUA activity was detected in various tissues 12 weeks after injection. (D) Tissue GAGs storage was detected in various tissues 12 weeks after injection. Wild-type C57/BL6 mice (n = 6) and Untreated *Idua-*W392X mice (n=6) were included as control. Treated *Idua*-W392X mice (n=5). Mean ± SD are shown. The treated *Idua-*W392X mice were compared with the untreated *Idua*-W392X mice, *p<0.05, **p<0.01, ***p<0.001, ****p<0.0001, Dunnett’s test.

In addition, we sacrificed the mice at 12 weeks post injection and harvested tissues to evaluate the tissue IDUA enzyme activity and GAGs storage. IDUA activity assay results showed that the IDUA activity of the heart and liver are significantly increased in the treated mice, reaching up to 27.3% and 17.3% of the activity in the wild-type mice, respectively. Slight increases of the IDUA activity in other tissues were also observed, corresponding to about 0.12% (spleen), 0.86% (lung), 1.65% (kidney), 1.3% (brain) of the activity in the wild-type mice, respectively, but with no significant difference (Figure 3C). Furthermore, the GAGs storage in peripheral tissues of the treated group were significantly reduced compared with the untreated MPS I mice, and there was no significant difference of the GAGs storage in heart and liver between the treated and the wild-type mice (Figure 3D).

### *In vivo* base editing reverses lysosomal storage damage in newborn MPS I mice

The accumulation of GAGs in tissues leads to the formation of characteristic microscopic lysosomal vacuoles (25). We performed a histological analysis of a subset of tissues (heart, liver, spleen, lung, kidney, and brain). In the hematoxylin and eosin (H&E) staining results, significant reduction of vacuolar cells was detected in the heart and liver tissues of the treated mice, and improvement in vacuolation of Purkinje cells was also observed (Figure 4A). A partial reduction of vacuolar cells was also observed in kidney and spleen tissues (Supplemental Figure 2). To evaluate correction of storage pathology in the treated mice, Alcian blue staining for GAGs was performed on tissue sections. Consistent with H&E staining results, variably decreased GAGs storage was observed in the heart, liver and brain tissues of treated mice (Figure 4B). In addition, histochemical analysis also showed no signs of inflammation such as lymphocyte or macrophage aggregation in tissues of the treated mice.

**Figure 4.**
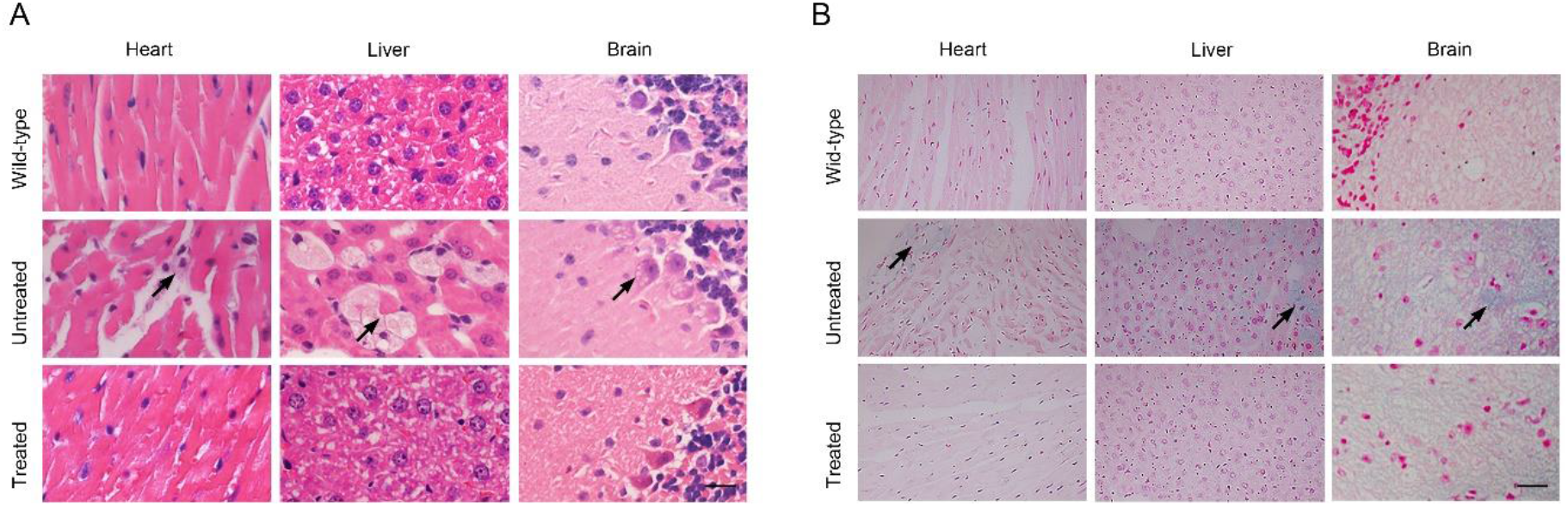
*In vivo* base editing corrects histological abnormalities in newborn MPS I mice. (A) Histological analysis of the heart, liver and brain at 12 weeks post-injection by hematoxylin and eosin stain. Scale bar, 20 μm. Black arrows indicate foamy macrophages in the tissue due to GAG accumulation. (B) The tissues were stained with Alcian blue to detect GAG. Scale bar, 20 μm. Black arrows indicate the GAGs storage in the tissues.

### *In vivo* base editing prevents neurobehavioral deficit in newborn MPS I mice

To detect whether base editors delivered *via* AAV9 provided any cognitive benefit to newborn MPS I mice, we performed a delayed-matching-to-place (DMP) dry maze test 12 weeks after injection. The DMP dry maze is a test that evaluates the learning and memory abilities of mice by measuring the time it takes for the mice to find an escape route on a high platform (26). After 4 days of testing and training, the average escape latency of wild-type mice with normal cognitive functions was reduced from 177s to 89s. In contrast, untreated MPS I mice showed a slow reduction in average escape latency from 170s to 138s, indicating cognitive deficits. Surprisingly, the escape of the treated mice was significantly faster on day 3 of the test, with no significant difference compared with the wild-type mice (Figure 5). The unexpected results suggested that *in viv*o base editing can effectively prevent cognitive deficits in newborn MPSI mice.

**Figure 5.**
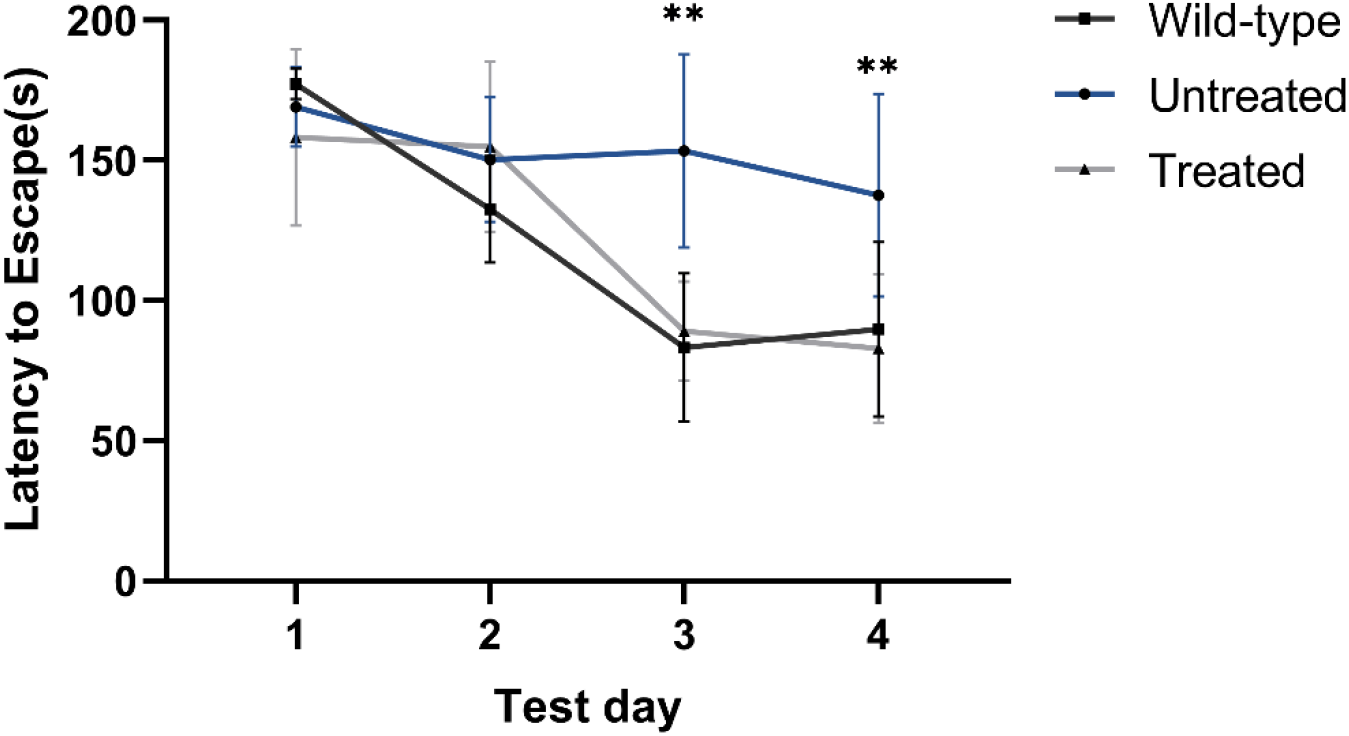
*In vivo* base editing prevents neurobehavioral deficit in newborn MPS I mice. Performance in the DMP dry maze is the time to escape from the maze. Data were shown as mean ±SD at each time point, for untreated *Idua*-W392X mice (n=6) compared with treated *Idua-*W392X mice (n=5). Wild-type C57/BL6 mice (n = 6) were also included as control. **p<0.01, Dunnett’s test.

## Discussion

There is no or limited treatment options for rare disease patients around the world, most of whom are suffering from monogenic diseases caused by single-nucleotide variants (SNV) (27). Base editing has the potential to correct SNV and can provide efficient and safe one-time treatment for many rare diseases (18). Herein, we demonstrated that AAV9-mediated split-intein ABE could effectively correct pathogenic mutations in newborn MPS I mice. We observed sustained serum IDUA activity and decreased tissue GAGs storage in MPS I treated mice. Moreover, the neurobehavioral deficits were significantly prevented.

MPS I is a multi-system disease involving the cardiovascular, respiratory, gastrointestinal, and nervous systems (28). ERT is the most extensive used treatment in the attenuated forms of MPS I, but is not recommended for the severe Hurler phenotype because the enzyme cannot cross the blood-brain barrier to influence the central nervous manifestations and cannot completely correct heart valvular or bone disease (29, 30). Despite early HSCT treatment may be able to prevent progressive neurocognitive impairment, the transplanted patients may still have a serious disease burden (31). Therefore, it is necessary to find a safer and more effective method to treat MPS I disease. An important feature of Mucopolysaccharidoses (MPSs) is its relatively low therapeutic threshold, which is extremely beneficial for the development of gene therapy/gene-editing therapies for these diseases (32, 33). In order to effectively treat central nervous manifestations of MPS I and prevent anti-transgenic immune response, Hinderer et al. performed systemic transgenic treatment of neonates before intrathecal administration, which effectively treated brain storage lesions (13). Vector dilution is a major problem in AAV gene therapy, which may lead to a gradual decline of therapeutic effect as the children grows (34). By contrast, AAV-mediated base editors can irreversibly correct the pathogenic genes and have a sustained therapeutic effect. In this study, we observed a high corrective efficiency in heart and liver tissues and improved disease outcomes 12 weeks after injection (Figure 2B). Additionally, we found that although the efficiency of genomic DNA correction in other tissues is low, the storage of GAGs is also reduced (Figure 3D). One possible explanation is that MPS I is a disease with a relatively low threshold for treatment (33). The second possibility is that the IDUA enzyme expressed and secreted in the heart and liver tissues is transmitted through the blood to other tissues, thereby reducing the GAGs storage in these tissues. Surprisingly, the prevention of neurobehavioral deficits was detected in the treated MPS I mice. We found a low vector copy numbers in the brain tissue, with a correction efficiency of about 0.34±0.12% (Figure 2B and 2C). Approximately 1.3% of the wild-type level of IDUA activity in the brain was observed in the treated mice (Figure 3C). It is worth noting that only 0.5% of wild-type activity is required to prevent neurological complications of MPS I (15).

In recent years, studies have reported that genome editing-mediated gene therapy could effectively repair the peripheral tissues and brain tissues of MPS I/MPS II mice (16, 35). However, the production of high frequency indels limits its clinical application. Previous studies showed that base editors did not randomly induce untargeted base conversion throughout the genome, but might cause unexpected editing in the regions where the sgRNA/base editor complex binds to DNA due to sequence homology (36-38). In this study, we estimated the top 10 potential off-target sites identified by a computer algorithm. NGS revealed that indels were less than 0.2% in highly edited liver tissues, suggesting that our base editing strategy is safer in MPS I treatment (Figure 2D). A recent study of intrauterine base editing in the treatment of MPS I mice has been reported, further confirming the effectiveness of base editing in MPS I (39). For progressive diseases such as MPS I, early treatment is more helpful to improve the disease outcomes. Many countries have introduced screening for neonatal lysosomal storage diseases. However, this screening is complicated by the wide clinical variability of these diseases and the fact that many people who are tested for enzyme deficiency will exhibit symptoms late or never in their lifetime (40). In addition, the operation of intrauterine injection therapy is difficult and risky, and requires very professional experts and equipment. In our research, the therapeutic effect on newborn mice is significant, and the operation is simple and has the potential for clinical application.

In conclusion, our results suggest that AAV-mediated base editor delivery can effectively correct storage damage in multiple tissues of genetic metabolic disease MPS I and prevent neurobehavioral deficits. Currently, there are many optimized ABE variants that are not only more efficient for editing but are no longer actually restricted by the requirement of PAM for sequence recognition. We believe that ABEs will become a favorable treatment for more genetic diseases caused by pathogenic mutations in the future.

## Material and Methods

### Plasmid Construction

VRQR-ABEmax (#119811), xCas9(3.7)-ABE(7.10) (#108382), NG-ABEmax (#124163), ABEmax(7.10)-SpG (#140002), NG-ABE8e (#138491) and pSPgRNA (#47108) plasmids were purchased from Addgene (Watertown, MA). To generate the ABE8e-SpG plasmid, ABE8e was digested by *NotI* and *EcoRV* and subcloned into ABEmax(7.10)-SpG plasmid backbone by In-Fusion cloning (Takara Bio, Mountain View, CA). The HEK293-*Idua* mutant cell lines were generated by stably integrating *Idua*-W392X sequence into the AAVS1 locus. CRISPR/Cas9 plasmid used to generate the HEK293-*Idua* mutant cell lines was constructed using pX330 (Plasmid #42230) (Supplemental Table 3). sgRNA-A5 and sgRNA-A6 targeting the G→A W392X mutation site on exon 9 of *Idua* gene in the mouse genome were designed by online webtool (https://benchling.com). All sgRNAs constructed were generated by T4 ligation of annealed oligos into *BbsI* digested pSPgRNA plasmid. Next, six adenine base editors (VRQR-ABEmax, xCas9(3.7)-ABE (x7.10), NG-ABEmax, ABEmax(7.10)-SpG, NG-ABE8e and ABE8e-SpG) were co-transfected with sgRNA-A5 or sgRNA-A6 into HEK293-*Idua* mutant cell lines, respectively. Genomic DNA was extracted 72h after transfection and Sanger sequencing was performed to screen the most effective base editors. The sgRNA-A6 was selected to further engineering of the split-intein dual-AAV system (referred to as N-ABE8e.SpG and C-ABE8e.SpG.sgRNA-A6). Both N-ABE8e.SpG and C-ABE8e.SpG.sgRNA-A6 vectors used CBh promoter and were generated by In-Fusion cloning of PCR-amplified inserting into restriction enzyme-digested backbones. All constructed plasmids were verified by sequencing.

### AAV Vector Production

AAV9.C-ABE8e-SpG and AAV9.N-ABE8e-SpG were obtained by packaging N-ABE8e.SpG and C-ABE8e.SpG.sgRNA-A6 into an AAV9 vectors (Supplemental Sequences). All AAV9 vectors were produced by triple plasmid transfection of HEK293 cells (ATCC, Manassas, VA) as previously described (41). The genome titer (genome copies [GCs] per milliliter, GC/ml) of AAV9 vector was determined by quantitative PCR (qPCR) using forward primer 5’-GCCAGCCATCTGTTGT-3’, reverse primer 5’-GGAGTGGCACCTTCCA-3’, and probe 5’-Fam-TCCCCCGTGCCTTCCTTGACC-Tamra-3’ (42). All vectors used in this study passed the endotoxin assay using the QCL-1000 Chromogenic LAL test kit (Cambrex Bio Science).

### Western blot analysis

Western blot analyses were performed on cell lysates. SpCas9 protein was detected by Mouse anti-CRISPR-Cas9 antibody (1:1000 dilution, Abcam, Cat# 191468). Mouse anti-GAPDH antibody (1:10000 dilution, ABclonal, Cat# AC002) was used to detect GAPDH. Blots were imaged and analyzed by iBrightTM CL1000 imaging systems (Thermo FisherScientific, InvitrogenTM).

### Animal studies

MPS I mice (*Idua*-W392X, Stock No: 017681) were purchased from Jackson Laboratory (Bar Harbor, Maine). The background of the wild-type mice used in this study were C57BL/6J. All animal protocols were approved by the Institutional Animal Care and Concern Committee at Sichuan University, and animal care was in accordance with the committee’s guidelines. Mating cages were monitored daily for births. Newborn (postnatal day 2, p2) pups received a temporal vein injection of the mixture of AAV9.C-SpG8e-SpG and AAV9.N-SpG8e-SpG at 1:1 (3×10 ^11^GC /mouse for each vector) in a volume of 50 μl, as described (43). Untreated wild-type, MPS I heterozygous (Het), and MPS I mice (*Idua*-W392X) served as controls. Mice were genotyped at weaning to confirm genotype. Serum samples for IDUA enzyme activity assays were obtained by retro-orbital bleeding 4 weeks post vector treatment and every 1 to 2 weeks thereafter. Urine samples were collected by gently applying pressure to the urinary bladder at the time of necropsy. The mice were killed at 12 weeks of age and tissues were collected for various analysis.

### IDUA enzyme activity assay

Tissue and serum samples were immediately frozen on dry ice and stored at -80° until analysis. Serum was used directly in IDUA enzyme activity assays. Tissue samples were homogenized in lysis buffer (0.9% NaCl, 0.2 % Triton-X100, pH 3.5), freeze-thawed and clarified by centrifugation. Protein concentrations were determined by BCA protein assay (Thermo Scientific, Waltham, MA). IDUA enzyme activity was determined in a fluorometric assay using the synthetic substrate 4MU-iduronide (Glycosynth, Warrington, England) as previously described (14). Units are given as nmol 4MU liberated per hour per mg of protein (tissues) or per ml of serum.

### Tissue GAGs assay

Tissue samples were consistent with IDUA enzyme assays. Tissue GAGs were determined using the Blyscan Glycosaminoglycan Assay Kit (Biocolor, Carrickfergus, UK), according to the manufacturer’s instructions.

### AAV9 biodistribution

DNA was extracted from tissues and total vector genomes quantified by Taqman qPCR as previously described (43).

### On-target and off-target analysis

To evaluate the on-target editing efficiency of various tissues, the tissue genomic DNA was extracted and then amplified by nest PCR to obtain the sequence fragment containing the W392X mutation, which was then analyzed by NGS. Furthermore, the top 10 potential off-target sites for sgRNA6 were identified by the algorithm described in www.benchling.com (Supplemental Table1). These off-target sites were amplified by nest PCR in the liver tissue genomic DNA and deep sequenced with NGS. Libraries were made from the second PCR products and sequenced on Illumina Miseq (2 × 300bp paired end, Personal Biotechnology Co., Ltd, Shanghai, China). Data were processed according to standard Illumina sequencing analysis procedures. Processed reads were mapped to the expected PCR amplicons as reference sequences using custom scripts. Reads that did not map to reference were discarded. Indels were determined by comparison of reads against reference using custom scripts.

### H&E staining

Tissues were fixed in paraformaldehyde for 24h, dehydrated through an ethanol series and xylene, and then embedded in paraffin. H&E staining was performed on 6 μm sections from paraffin-embedded tissues according to standard protocols.

### GAGs histochemistry

Tissue samples were prepared as H&E staining. Deparaffinized 6 μm sections were stained in 1% Alcian Blue (Sigma, #MKCM1030) for 15 minutes, rinsed in water for 2–3 minutes, and counterstained with Nuclear Fast Red (Sigma, #N8002).

### DMP dry maze assay

To detect whether base editors delivered *via* AAV9 provided any cognitive benefit to MPS I mice, we performed a DMP dry maze test 12 weeks after injection. DMP dry maze test was a variant of DMP water maze (44). The DMP dry maze was a circular platform (Diameter = 122 cm, thickness = 1.2 cm) with 40 holes. An escape pipe was secured under one of the holes to allow the mice to escape the platform. The location of the escape hole changed every day. Visual cues were attached to each of the four walls for the mouse to use in spatial navigation. To begin the experiment, mice were placed on the edge of a platform some distance from the escape hole, and an opaque funnel covered the mouse. After a delay of about 30 sec, turning on the tone noise (2 KHz, 85 dB) and immediately removing the transfer box to expose mice in a bright light (1200 Lux). In response to these aversive conditions, the mice would spontaneously seek out and burrow into the escape hole. Mice were assessed during four trials per day on four consecutive days, with a maximal escape time limited to 3 min. Data were collected and analyzed using the ANY-Maze program.

### Statistics

Graphpad Prism9 was used to perform all statistical tests. Values express mean ± SD. Statistical analysis was by Dunnett’s test, as indicated in figure legends. In all tests, p<0.05 was considered significant.

## Author Contributions

Y.Y. conceived this study and designed the experiments; Y.L. constructed the plasmid vectors; X.J. produced AAV9 vector and endotoxin assays; J.S. and X.J. performed mouse studies; J.S. performed on and off-target analyses; K.S., X.Z. and J.X. performed DMP dry maze assay; Q.Z. performed qPCR analysis; J.S. and R.L. performed histopathology assays; J.S. wrote the manuscript; Y.Y. and H.D. edited the manuscript. All authors read and approved the final manuscript.

## Acknowledgments

This work was supported by the Joint Funds of the National Natural Science Foundation of China (Grant No.U19A2002), National Major Scientific and Technological Special Project for “Significant New Drugs Development” (No.2018ZX09733001-005-002), and the Science and Technology Major Project of Sichuan province (No.2017SZDZX0011).

## References

1. Coutinho MF, Lacerda L, and Alves S. Glycosaminoglycan storage disorders: a review. Biochem Res Int. 2012;2012:471325.

2. Hampe CS, Eisengart JB, Lund TC, Orchard PJ, Swietlicka M, Wesley J, et al. Mucopolysaccharidosis Type I: A Review of the Natural History and Molecular Pathology. Cells. 2020;9(8).

3. Moore D, Connock MJ, Wraith E, and Lavery C. The prevalence of and survival in Mucopolysaccharidosis I: Hurler, Hurler-Scheie and Scheie syndromes in the UK. Orphanet J Rare Dis. 2008;3:24.

4. Tebani A, Zanoutene-Cheriet L, Adjtoutah Z, Abily-Donval L, Brasse-Lagnel C, Laquerriere A, et al. Clinical and Molecular Characterization of Patients with Mucopolysaccharidosis Type I in an Algerian Series. Int J Mol Sci. 2016;17(5).

5. Thomas S, and Tandon S. Hurler syndrome: a case report. Journal of Clinical Pediatric Dentistry. 2008;24(4):335–8.

6. Scott HS, Litjens T, Hopwood JJ, and Morris CP. A common mutation for mucopolysaccharidosis type I associated with a severe Hurler syndrome phenotype. Hum Mutat. 1992;1(2):103–8.

7. Pineda T, Marie S, Gonzalez J, Garcia AL, Acosta A, Morales M, et al. Genotypic and bioinformatics evaluation of the alpha-l-iduronidase gene and protein in patients with mucopolysaccharidosis type I from Colombia, Ecuador and Peru. Mol Genet Metab Rep. 2014;1:468–73.

8. Poletto E, Pasqualim G, Giugliani R, Matte U, and Baldo G. Worldwide distribution of common IDUA pathogenic variants. Clinical Genetics. 2018;94(1):95–102.

9. Clarke LA, Atherton AM, Burton BK, Day-Salvatore DL, Kaplan P, Leslie ND, et al. Mucopolysaccharidosis Type I Newborn Screening: Best Practices for Diagnosis and Management. J Pediatr. 2017;182:363–70.

10. Tolar J, Grewal SS, Bjoraker KJ, Whitley CB, Shapiro EG, Charnas L, et al. Combination of enzyme replacement and hematopoietic stem cell transplantation as therapy for Hurler syndrome. Bone Marrow Transplant. 2008;41(6):531–5.

11. Parini R, Deodato F, Di Rocco M, Lanino E, Locatelli F, Messina C, et al. Open issues in Mucopolysaccharidosis type I-Hurler. Orphanet J Rare Dis. 2017;12(1):112.

12. Hinderer C, Bell P, Gurda BL, Wang Q, Louboutin J-P, Zhu Y, et al. Liver-directed gene therapy corrects cardiovascular lesions in feline mucopolysaccharidosis type I. Proceedings of the National Academy of Sciences. 2014;111(41):14894–9.

13. Hinderer C, Bell P, Louboutin JP, Zhu Y, Yu H, Lin G, et al. Neonatal Systemic AAV Induces Tolerance to CNS Gene Therapy in MPS I Dogs and Nonhuman Primates. Mol Ther. 2015;23(8):1298–307.

14. Hinderer C, Bell P, Gurda BL, Wang Q, Louboutin JP, Zhu Y, et al. Intrathecal gene therapy corrects CNS pathology in a feline model of mucopolysaccharidosis I. Mol Ther. 2014;22(12):2018–27.

15. Ou L, Przybilla MJ, Ahlat O, Kim S, Overn P, Jarnes J, et al. A Highly Efficacious PS Gene Editing System Corrects Metabolic and Neurological Complications of Mucopolysaccharidosis Type I. Mol Ther. 2020;28(6):1442–54.

16. Ou L, DeKelver RC, Rohde M, Tom S, Radeke R, St Martin SJ, et al. ZFN-Mediated In Vivo Genome Editing Corrects Murine Hurler Syndrome. Mol Ther. 2019;27(1):178–87.

17. Rees HA, and Liu DR. Base editing: precision chemistry on the genome and transcriptome of living cells. Nat Rev Genet. 2018;19(12):770–88.

18. Porto EM, Komor AC, Slaymaker IM, and Yeo GW. Base editing: advances and therapeutic opportunities. Nat Rev Drug Discov. 2020;19(12):839–59.

19. Wang D, Shukla C, Liu X, Schoeb TR, Clarke LA, Bedwell DM, et al. Characterization of an MPS I-H knock-in mouse that carries a nonsense mutation analogous to the human IDUA-W402X mutation. Mol Genet Metab. 2010;99(1):62–71.

20. Richter MF, Zhao KT, Eton E, Lapinaite A, Newby GA, Thuronyi BW, et al. Phage-assisted evolution of an adenine base editor with improved Cas domain compatibility and activity. Nature Biotechnology. 2020;38(7):883–91.

21. Huang TP, Zhao KT, Miller SM, Gaudelli NM, Oakes BL, Fellmann C, et al. Circularly permuted and PAM-modified Cas9 variants broaden the targeting scope of base editors. Nature Biotechnology. 2019;37(6):626–31.

22. Zincarelli C, Soltys S, Rengo G, and Rabinowitz JE. Analysis of AAV serotypes 1-9 mediated gene expression and tropism in mice after systemic injection. Mol Ther. 2008;16(6):1073–80.

23. Inagaki K, Fuess S, Storm TA, Gibson GA, McTiernan CF, Kay MA, et al. Robust systemic transduction with AAV9 vectors in mice: efficient global cardiac gene transfer superior to that of AAV8. Mol Ther. 2006;14(1):45–53.

24. Kiely BT, Kohler JL, Coletti HY, Poe MD, and Escolar ML. Early disease progression of Hurler syndrome. Orphanet J Rare Dis. 2017;12(1):32.

25. Ohmi K, Greenberg DS, Rajavel KS, Ryazantsev S, Li HH, and Neufeld EF. Activated microglia in cortex of mouse models of mucopolysaccharidoses I and IIIB. Proc Natl Acad Sci U S A. 2003;100(4):1902–7.

26. Feng X, Krukowski K, Jopson T, and Rosi S. Delayed-matching-to-place Task in a Dry Maze to Measure Spatial Working Memory in Mice. Bio Protoc. 2017;7(13).

27. Posey JE. Genome sequencing and implications for rare disorders. Orphanet Journal of Rare Diseases. 2019;14(1).

28. Kubaski F, de Oliveira Poswar F, Michelin-Tirelli K, Matte UdS, Horovitz DD, Barth AL, et al. Mucopolysaccharidosis Type I. Diagnostics. 2020;10(3):161.

29. Miebach E. Enzyme replacement therapy in mucopolysaccharidosis type I. Acta Paediatrica. 2005;94(447):58–60.

30. Wraith JE. Enzyme replacement therapy in mucopolysaccharidosis type I: progress and emerging difficulties. J Inherit Metab Dis. 2001;24(2):245–50.

31. Hampe CS, Wesley J, Lund TC, Orchard PJ, Polgreen LE, Eisengart JB, et al. Mucopolysaccharidosis Type I: Current Treatments, Limitations, and Prospects for Improvement. Biomolecules. 2021;11(2).

32. Sands MS, and Davidson BL. Gene therapy for lysosomal storage diseases. Mol Ther. 2006;13(5):839–49.

33. Poletto E, Baldo G, and Gomez-Ospina N. Genome Editing for Mucopolysaccharidoses. Int J Mol Sci. 2020;21(2).

34. Verdera HC, Kuranda K, and Mingozzi F. AAV Vector Immunogenicity in Humans: A Long Journey to Successful Gene Transfer. Mol Ther. 2020;28(3):723–46.

35. Laoharawee K, DeKelver RC, Podetz-Pedersen KM, Rohde M, Sproul S, Nguyen HO, et al. Dose-Dependent Prevention of Metabolic and Neurologic Disease in Murine MPS II by ZFN-Mediated In Vivo Genome Editing. Mol Ther. 2018;26(4):1127–36.

36. Villiger L, Grisch-Chan HM, Lindsay H, Ringnalda F, Pogliano CB, Allegri G, et al. Treatment of a metabolic liver disease by in vivo genome base editing in adult mice. Nature Medicine. 2018;24(10):1519–25.

37. Suh S, Choi EH, Leinonen H, Foik AT, Newby GA, Yeh W-H, et al. Restoration of visual function in adult mice with an inherited retinal disease via adenine base editing. Nature Biomedical Engineering. 2021;5(2):169–78.

38. Koblan LW, Erdos MR, Wilson C, Cabral WA, Levy JM, Xiong Z-M, et al. In vivo base editing rescues Hutchinson–Gilford progeria syndrome in mice. Nature. 2021;589(7843):608–14.

39. Bose SK, White BM, Kashyap MV, Dave A, De Bie FR, Li H, et al. In utero adenine base editing corrects multi-organ pathology in a lethal lysosomal storage disease. Nat Commun. 2021;12(1):4291.

40. Beck M. Treatment strategies for lysosomal storage disorders. Developmental Medicine & Child Neurology. 2018;60(1):13–8.

41. Lock M, Alvira M, Vandenberghe LH, Samanta A, Toelen J, Debyser Z, et al. Rapid, simple, and versatile manufacturing of recombinant adeno-associated viral vectors at scale. Hum Gene Ther. 2010;21(10):1259–71.

42. Lock M, Alvira MR, Chen SJ, and Wilson JM. Absolute determination of single-stranded and self-complementary adeno-associated viral vector genome titers by droplet digital PCR. Hum Gene Ther Methods. 2014;25(2):115–25.

43. Yang Y, Wang L, Bell P, McMenamin D, He Z, White J, et al. A dual AAV system enables the Cas9-mediated correction of a metabolic liver disease in newborn mice. Nature biotechnology. 2016;34(3):334–8.

44. Faizi M, Bader PL, Saw N, Nguyen TV, Beraki S, Wyss-Coray T, et al. Thy1-hAPP(Lond/Swe+) mouse model of Alzheimer’s disease displays broad behavioral deficits in sensorimotor, cognitive and social function. Brain Behav. 2012;2(2):142–54.

