## Supplemental material for "*In vivo* adenine base editing corrects newborn murine model of Hurler syndrome"

*Corresponding authors: Yang Yang

Tel: + 86 028 85164063

### Supplemental Figure 1

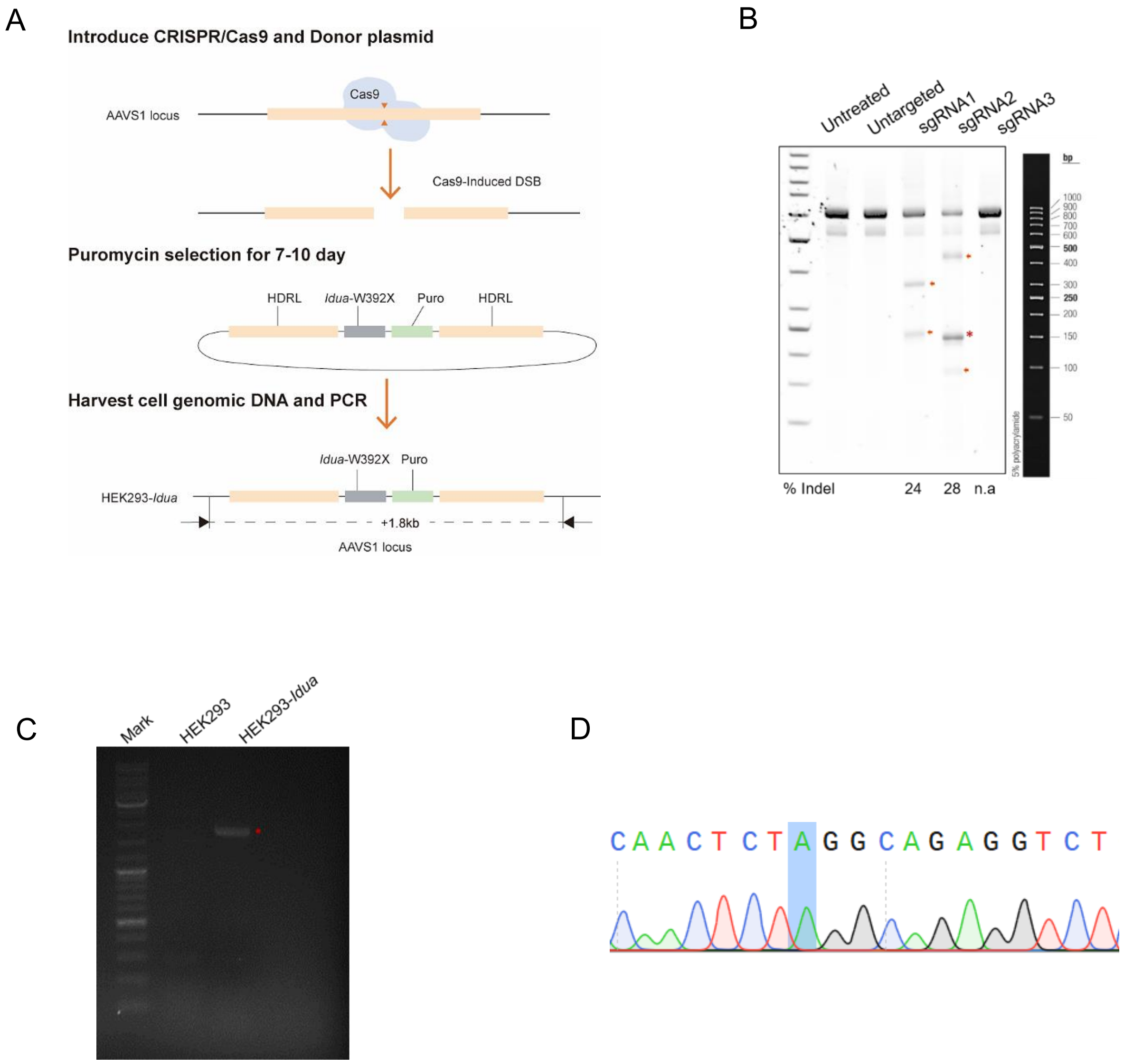


**Fig.S1. Construction of HEK293-*Idua* mutant cell line using CRISPR/Cas9. (A)** Schematic diagram of HEK293-*Idua* mutant cell line construction. (B) Screening of sgRNA in the construction of mutant cell lines. *In vitro* validation of the editing effect of sgRNAs in the HEK293 cell line by transient transfection and SURVEYOR nuclease assays. Arrows denote SURVEYOR nuclease cleaved fragments of the AAVS1 PCR products. Asterisks indicate nonspecific bands. (C) Gel electrophoresis verified that the mutant sequence was successfully inserted into the DNA genome of HEK293 cells. The inserted mutant sequences are marked in red (D) The successful construction of HEKK293-*Idua* mutant cell line was verified by Sanger sequencing. The shaded part is the mutation site.

**Supplemental Figure 2**

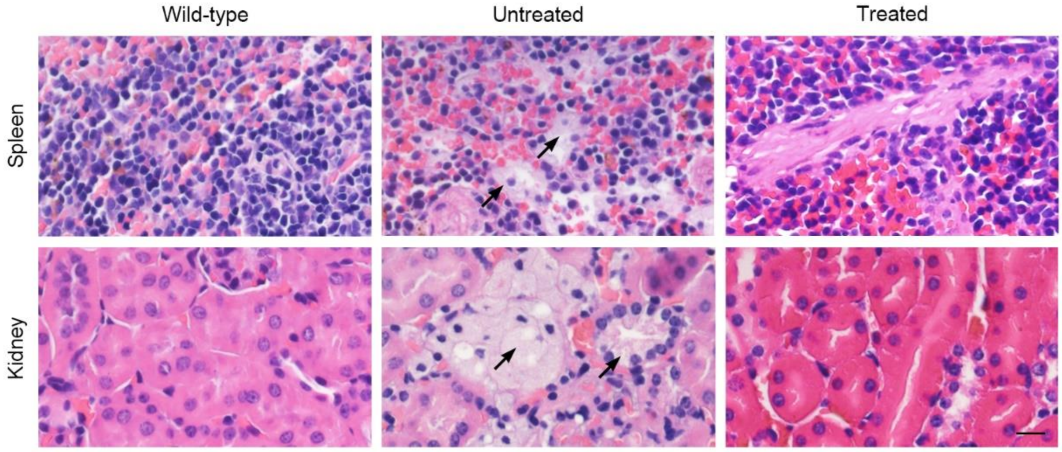


**Fig.S2. In vivo base editing can correct histological abnormalities in MPSI mice.** Histological analysis of the spleen and kidney at 12 weeks post-injection by hematoxylin and eosin stain. Scale bar, 20μm. Black arrows indicate foamy macrophages in the tissue due to GAG accumulation.

**Supplemental Table 1. Primers and sequences for construction of HEK293-*Idua* mutant cell lines.**

| **Name** | **Sequence** | **Note** |
|---|---|---|
| sgRNA1 | GGGACCACCTTATATTCCCA (PAM: GGG) | Three sgRNA sequences for construction of HEK293-*Idua* mutant cell lines |
| sgRNA2 | GAGATGGCTCCAGGAAATGG (PAM: GGG) | |
| sgRNA3 | TAAGGAATCTGCCTAACAGG (PAM: AGG) | |
| AAVS1_T1 Fwd | CACCGGGACCACCTTATATTCCCA | CRISPR/Cas9 plasmid-1 construction using pX330 |
| AAVS1_T1 Rev | AAACTGGGAATATAAGGTGGTCCC | |
| AAVS1_T2Fwd | CACCGAGATGGCTCCAGGAAATGG | CRISPR/Cas9 plasmid-2 construction using pX330 |
| AAVS1_T2 Rev | AAACCCATTTCCTGGAGCCATCTC | |
| AAVS1_T3 Fwd | CACCGTAAGGAATCTGCCTAACAGG | CRISPR/Cas9 plasmid-3 construction using pX330 |
| AAVS1_T3 Rev | AAACCCTGTTAGGCAGATTCCTTAC | |
| AAVS1_ P1Fwd | TCCTGAGTCCGGACCACTTT | The primers of surveyor assay for validation of sgRNA |
| AAVS1_ P1Rev | GCTTCTTGGCCACGTAACCT | |
| AAVS1_ PointMF | TTTGCCTGGACACCCCGTTC | |
| AAVS1_ PointMR | GCGTCAGAGCAGCTCAGGTT | |
| MPS1_Fwd | CCTGGCACATCCTGTATTGA | PCR Primers for HEK293-*Idua* mutant cell lines |
| MPS1_Rev | CCTCTACAAATGTGGTATGGC | |

### Supplemental Table 2. Off-target analysis. Potential off-target sequences for sgRNA-A6 identified and scored by Benchling's off-target analysis.

| ID | Sequence | PAM | Score | Chromosome | Strand | Position | Mismatches | On-target |
|---|---|---|---|---|---|---|---|---|
| **SgRNA-A6** | ACTCTGGGCAGAGGTCTCAA | AG | 100.00 | Chr5 | 1 | 108681407 | 0 | |
| **OT1** | TGGCTAGGCAGAGGTCTCAA | TG | 2.32 | Chr11 | -1 | 72424740 | 3 | FALSE |
| **OT2** | AATGTATGCAGAGGTCTCAA | GG | 1.77 | Chr6 | -1 | 76674529 | 3 | FALSE |
| **OT3** | TCTCTAGAAAGAGGTCTCAA | AG | 1.75 | Chr8 | -1 | 34402855 | 3 | FALSE |
| **OT4** | ACTTTATGCTGAGGTCTCAA | TG | 1.68 | Chr7 | -1 | 64896671 | 3 | FALSE |
| **OT5** | ACTCCAGCAAGAGGTCTCAA | AG | 1.53 | chr12 | -1 | 81903353 | 3 | FALSE |
| **OT6** | GTTCTAGACTGAGGTCTCAA | GG | 1.50 | Chr17 | -1 | 8999275 | 4 | FALSE |
| **OT7** | AGTTCAGACAGAGGTCTCAA | AG | 1.44 | Chr9 | -1 | 26721248 | 4 | FALSE |
| **OT8** | ATCCAAGGCTGAGGTCTCAA | TG | 1.39 | chr17 | 1 | 33748567 | 4 | FALSE |
| **OT9** | TTACAAGGCAGAGGTCTCAA | TG | 1.36 | chr11 | -1 | 111283516 | 4 | FALSE |
| **OT10** | TTCCCAGGCAGAGGTCTCAA | AG | 1.36 | Chr8 | 1 | 69133495 | 4 | FALSE |

**Supplemental Table 3. PCR primer sequences for detecting potential on-target and off-target effects by NGS assay.**

| Primer Name | Sequence | Note |
|---|---|---|
| **On-target** | | |
| Nest_P1Fwd | AGTTGCTGCGAAAGCCAGTA | Primers for on-target |
| Nest1_P1Rev | AGGGATACTGTGGTTGGGGT | |
| Nest2_P2FWD | GGTGGGAGCTAGATATTAGG | |
| Nest2_P2Rev | AGATGAGGACTGTGGTACTC | |
| **Off-target** | | |
| OT1_P1Fwd | ACTGGAAGAAACAGCGGAAG | Primers for off-target1 |
| OT1_P1Rev | GAGTGTGAGACCTCTAAGAG | |
| OT1_P2Fwd | AGCTGGGGAACAAGTGTTCA | |
| OT1_P2Rev | GCTGGATAACTACACCGTTC | |
| OT2_P1Fwd | CACTGGTTTCCAAGACTAAGG | Primers for off-target2 |
| OT2_P1Rev | AGGTTAAGTGCAGAAAGGAG | |
| OT2_P2Fwd | ACCAATACAGTAGCCCTATG | |
| OT2_P2Rev | GCAGAAAGGAGATATTTCTCTGAG | |
| OT3_P1Fwd | CTCTAGCTCTCATGCACACA | Primers for off-target3 |
| OT3_P1Rev | CCCTCAGTGGCAACCTATAA | |
| OT3_P2Fwd | TTGGAGACCTGTGACTAGGA | |
| OT3_P2Rev | TGGGTCACTTGAGTTCTTCC | |
| OT4_P1Fwd | CCGTTGTGAGTGTGCCATTT | Primers for off-target4 |
| OT4_P1Rev | GATCTGTCCACTGATCCACT | |
| OT4_P2Fwd | GACCAAACAAGGACTTGTCA | |
| OT4_P2Rev | GCACACTTAGGTCTACCTTC | |
| OT5_P1Fwd | GGCAAGACAATCTTGCTCAC | Primers for off-target5 |
| OT5_P1Rev | TACGTGGCAGTGAGGAATTG | |
| OT5_P2Fwd | GTGTTTCTGATGAAACTTCC | |
| OT5_P2Rev | AGGAGCAGATTCTCTCACAC | |
| OT6_P1Fwd | GAACTGAGAGGAGAGCGAAA | Primers for off-target6 |
| OT6_P1Rev | CCAGAAGGCTCTCTGCATTT | |
| OT6_P2Fwd | GTAAGAGTGGGCCATGTGAA | |
| OT6_P2Rev | ATTCTGGCAGGTGCTGCATGTA | |
| OT7_P1Fwd | GATCACACAATGGGCACTCT | Primers for off-target7 |
| OT7_P1Rev | GAGTGGTTGGAAGTTCAAGC | |
| OT7_P2Fwd | TTCTATCCTGTTGGGAGGCA | |
| OT7_P2Rev | ATAGGGTGTTGAGATTCTGG | |
| OT8_P1Fwd | GTTAGCTGCTTCTCCCTAGA | Primers for off-target8 |
| OT8_P1Rev | ACCTGAGTCACTGTTCCATC | |
| OT8_P2Fwd | CTGTGCTCTGTATCTGCACA | |
| OT8_P2Rev | ACAGTTGCATCCTGGAGCTA | |
| OT9_P1Fwd | AGGCTGCTTGTTCCTTCCTT | Primers for off-target9 |

| | | |
|---|---|---|
| OT9_P1Rev | CAGTGTCAAGGCTCATAGTC | |
| OT9_P2Fwd | ACTGGGTCCTACACATAAAG | |
| OT9_P2Rev | TGGGAGCCAGACCCTCTTATTT | |
| OT10_P1Fwd | TCCTACTCTCTTAACTCCC | Primers for off-target10 |
| OT10_P1Rev | TCTATTCAGTGTGCCCTGGT | |
| OT10_P2Fwd | CTTCCCCCATACCTGATTAT | |
| OT10_P2Rev | TGCCTCCAAGGAAAGACTCA | |

### Supplemental Sequences. Coding sequences of split-intein ABE

#### Coding sequence for N-ABE8e.SpG-Int$^N$

MKRTADGSEFESPKKKRKVSEVEFSHEYWMRHALTLAKRARDEREVPVGAVLVLNNRVIGEGWNRAIGL
HDPTAHAEIMALRQGGLVMQNYRLIDATLYVTFEPCVMCAGAMIHSRIGRVVFGVRNSKRGAAGSLMNV
LNYPGMNHRVEITEGILADECAALLCDFYRMPRQVFNAQKKAQSSINSGGSSGGSSGSETPGTSESATPES
SGGSSGGSDKKYSIGLAIGTNSVGWAVITDEYKVPSKKFKVLGNTDRHSIKKNLIGALLFDSGETAEATRL
KRTARRRYTRRKNRICYLQEIFSNEMAKVDDSFFHRLEESFLVEEDKKHERHPIFGNIVDEVAYHEKYPTIY
HLRKKLVDSTDKADLRLIYLALAHMIKFRGHFLIEGDLNPDNSDVDKLFIQLVQTYNQLFEENPINASGVD
AKAILSARLSKSRRLENLIAQLPGEKKNGLFGNLIALSLGLTPNFKSNFDLAEDAKLQLSKDTYDDDLDNL
LAQIGDQYADLFLAAKNLSDAILLSDILRVNTEITKAPLSASMIKRYDEHHQDLTLLKALVRQQLPEKYKEI
FFDQSKNGYAGYIDGGASQEEFYKFIKPILEKMDGTEELLVKLNREDLLRKQRTFDNGSIPHQIHLGELHAI
LRRQEDFYPFLKDNREKIEKILTFRIPYYVGPLARGNSRFAWMTRKSEETITPWNFEEVVDKGASAQSFIER
MTNFDKNLPNEKVLPKHSLLYEYFTVYNELTKVKYVTEGMRKPAFLSGEQKKAIVDLLFKTNRKVTVKQ
LKEDYFKKIECLSYETEILTVEYGLLPIGKIVEKRIECTVYSVDNNGNIYTQPVAQWHDRGEQEVFEYCLED
GSLIRATKDHKFMTVDGQMLPIDEIFERELDLMRVDNLPN

ATGAAACGGACAGCCGACGGAAGCGAGTTCGAGTCACCAAAGAAGAAGCGGAAAGTCTCTGAGGTG
GAGTTTTCCCACGAGTACTGGATGAGACATGCCCTGACCCTGGCCAAGAGGGCACGGGATGAGAGGG
AGGTGCCTGTGGGAGCCGTGCTGGTGCTGAACAATAGAGTGATCGGCGAGGGCTGGAACAGAGCCAT
CGGCCTGCACGACCCAACAGCCCATGCCGAAATTATGGCCCTGAGACAGGGCGGCCTGGTCATGCAG
AACTACAGACTGATTGACGCCACCCTGTACGTGACATTCGAGCCTTGCGTGATGTGCGCCGGCGCCAT
GATCCACTCTAGGATCGGCCGCGTGGTGTTTGGCGTGAGGAACTCAAAAAGAGGCGCCGCAGGCTCC
CTGATGAACGTGCTGAACTACCCCGGCATGAATCACCGCGTCGAAATTACCGAGGGAATCCTGGCAGA
TGAATGTGCCGCCCTGCTGTGCGATTTCTATCGGATGCCTAGACAGGTGTTCAATGCTCAGAAGAAGG
CCCAGAGCTCCATCAACTCCGGAGGATCTAGCGGAGGCTCCTCTGGCTCTGAGACACCTGGCACAAG
CGAGAGCGCAACACCTGAAAGCAGCGGGGGCAGCAGCGGGGGGTCAGACAAGAAGTACAGCATCGG
CCTGGCCATCGGCACCAACTCTGTGGGCTGGGCCGTGATCACCGACGAGTACAAGGTGCCCAGCAAG
AAATTCAAGGTGCTGGGCAACACCGACCGGCACAGCATCAAGAAGAACCTGATCGGAGCCCTGCTGT
TCGACAGCGGCGAAACAGCCGAGGCCACCCGGCTGAAGAGAACCGCCAGAAGAAGATACACCAGAC
GGAAGAACCGGATCTGCTATCTGCAAGAGATCTTCAGCAACGAGATGGCCAAGGTGGACGACAGCTT
CTTCCACAGACTGGAAGAGTCCTTCCTGGTGGAAGAGGATAAGAAGCACGAGCGGCACCCCATCTTC
GGCAACATCGTGGACGAGGTGGCCTACCACGAGAAGTACCCCACCATCTACCACCTGAGAAAGAAAC
TGGTGGACAGCACCGACAAGGCCGACCTGCGGCTGATCTATCTGGCCCTGGCCCACATGATCAAGTTC
CGGGGCCACTTCCTGATCGAGGGCGACCTGAACCCCGACAACAGCGACGTGGACAAGCTGTTCATCC
AGCTGGTGCAGACCTACAACCAGCTGTTCGAGGAAAACCCCATCAACGCCAGCGGCGTGGACGCCAA
GGCCATCCTGTCTGCCAGACTGAGCAAGAGCAGACGGCTGGAAAATCTGATCGCCCAGCTGCCCGGC
GAGAAGAAGAATGGCCTGTTCGGAAACCTGATTGCCCTGAGCCTGGGCCTGACCCCCAACTTCAAGA
GCAACTTCGACCTGGCCGAGGATGCCAAACTGCAGCTGAGCAAGGACACCTACGACGACGACCTGG
ACAACCTGCTGGCCCAGATCGGCGACCAGTACGCCGACCTGTTTCTGGCCGCCAAGAACCTGTCCGA
CGCCATCCTGCTGAGCGACATCCTGAGAGTGAACACCGAGATCACCAAGGCCCCCCTGAGCGCCTCT
ATGATCAAGAGATACGACGAGCACCACCAGGACCTGACCCTGCTGAAAGCTCTCGTGCGGCAGCAGC
TGCCTGAGAAGTACAAAGAGATTTTCTTCGACCAGAGCAAGAACGGCTACGCCGGCTACATTGACGG

CGGAGCCAGCCAGGAAGAGTTCTACAAGTTCATCAAGCCCATCCTGGAAAAGATGGACGGCACCGAG
GAACTGCTCGTGAAGCTGAACAGAGAGGACCTGCTGCGGAAGCAGCGGACCTTCGACAACGGCAGC
ATCCCCCACCAGATCCACCTGGGAGAGCTGCACGCCATTCTGCGGCGGCAGGAAGATTTTTACCCATT
CCTGAAGGACAACCGGGAAAAGATCGAGAAGATCCTGACCTTCCGCATCCCCTACTACGTGGGCCCTC
TGGCCAGGGGAAACAGCAGATTCGCCTGGATGACCAGAAAGAGCGAGGAAACCATCACCCCCTGGA
ACTTCGAGGAAGTGGTGGACAAGGGCGCTTCCGCCCAGAGCTTCATCGAGCGGATGACCAACTTCGA
TAAGAACCTGCCCAACGAGAAGGTGCTGCCCAAGCACAGCCTGCTGTACGAGTACTTCACCGTGTATA
ACGAGCTGACCAAAGTGAAATACGTGACCGAGGGAATGAGAAAGCCCGCCTTCCTGAGCGGCGAGC
AGAAAAAGGCCATCGTGGACCTGCTGTTCAAGACCAACCGGAAAGTGACCGTGAAGCAGCTGAAAG
AGGACTACTTCAAGAAAATCGAGTGCCTGAGCTACGAGACAGAGATCCTGACCGTGGAATACGGCCT
GCTGCCTATCGGCAAGATCGTGGAAAAGCGGATCGAGTGCACCGTGTACAGCGTGGACAACAACGGC
AACATCTACACCCAGCCTGTGGCTCAGTGGCACGACAGAGGCGAGCAAGAGGTGTTCGAGTACTGCC
TGGAAGATGGCAGCCTGATCAGAGCCACCAAGGACCACAAGTTCATGACAGTGGACGGCCAGATGCT
GCCCATCGACGAGATCTTCGAGCGCGAGCTGGACCTGATGAGAGTGGACAACCTGCCTAACTAA

### Coding sequence for Int$^{c}$-C-ABE8e.SpG

MIKIATRKYLGKQNVYDIGVERDHNFALKNGFIASNQSGKTILDFLKSDGFANRNFMQLIHDDSLTFKEDI
QKAQVSGQGDSLHEHIANLAGSPAIKKGILQTVKVVDELVKVMGRHKPENIVIEMARENQTTQKGQKNS
RERMKRIEEGIKELGSQILKEHPVENTQLQNEKLYLYYLQNGRDMYVDQELDINRLSDYDVDHIVPQSFL
KDDSIDNKVLTRSDKNRGKSDNVPSEEVVKKMKNYWRQLLNAKLITQRKFDNLTKAERGGLSELDKAGF
IKRQLVETRQITKHVAQILDSRMNTKYDENDKLIREVKVITLKSKLVSDFRKDFQFYKVREINNYHHAHDA
YLNAVVGTALIKKYPKLESEFVYGDYKVYDVRKMIAKSEQEIGKATAKYFFYSNIMNFFKTEITLANGEIR
KRPLIETNGETGEIVWDKGRDFATVRKVLSMPQVNIVKKTEVQTGGFSKESILPKRNSDKLIARKKDWDP
KKYGGFLWPTVAYSVLVVAKVEKGKSKKLKSVKELLGITIMERSSFEKNPIDFLEAKGYKEVKKDLIIKLP
KYSLFELENGRKRMLASAKQLQKGNELALPSKYVNFLYLASHYEKLKGSPEDNEQKQLFVEQHKHYLDE
IIEQISEFSKRVILADANLDKVLSAYNKHRDKPIREQAENIIHLFTLTNLGAPAAFKYFDTTIDRKQYRSTKE
VLDATLIHQSITGLYETRIDLSQLGGDSGGSKRTADGSEFEPKKKRKV

ATGATCAAGATCGCCACACGGAAGTACCTGGGCAAGCAGAACGTGTACGACATCGGCGTGGAACGGG
ACCACAACTTCGCCCTGAAGAACGGCTTTATCGCCAGCAACTGCTTCGACTCCGTGGAAATCTCCGGC
GTGGAAGATCGGTTCAACGCCTCCCTGGGCACATACCACGATCTGCTGAAAATTATCAAGGACAAGGA
CTTCCTGGACAATGAGGAAAACGAGGACATTCTGGAAGATATCGTGCTGACCCTGACACTGTTTGAGG
ACAGAGAGATGATCGAGGAACGGCTGAAAACCTATGCCCACCTGTTCGACGACAAAGTGATGAAGCA
GCTGAAGCGGCGGAGATACACCGGCTGGGGCAGGCTGAGCCGGAAGCTGATCAACGGCATCCGGGA
CAAGCAGTCCGGCAAGACAATCCTGGATTTCCTGAAGTCCGACGGCTTCGCCAACAGAAACTTCATG
CAGCTGATCCACGACGACAGCCTGACCTTTAAAGAGGACATCCAGAAAGCCCAGGTGTCCGGCCAGG
GCGATAGCCTGCACGAGCACATTGCCAATCTGGCCGGCAGCCCCGCCATTAAGAAGGGCATCCTGCAG
ACAGTGAAGGTGGTGGACGAGCTCGTGAAAGTGATGGGCCGGCACAAGCCCGAGAACATCGTGATC
GAAATGGCCAGAGAGAACCAGACCACCCAGAAGGGACAGAAGAACAGCCGCGAGAGAATGAAGCG
GATCGAAGAGGGCATCAAAGAGCTGGGCAGCCAGATCCTGAAAGAACACCCCGTGGAAAACACCCA

GCTGCAGAACGAGAAGCTGTACCTGTACTACCTGCAGAATGGGCGGGATATGTACGTGGACCAGGAA
CTGGACATCAACCGGCTGTCCGACTACGATGTGGACCATATCGTGCCTCAGAGCTTTCTGAAGGACGA
CTCCATCGACAACAAGGTGCTGACCAGAAGCGACAAGAACCGGGGCAAGAGCGACAACGTGCCCTC
CGAAGAGGTCGTGAAGAAGATGAAGAACTACTGGCGGCAGCTGCTGAACGCCAAGCTGATTACCCAG
AGAAAGTTCGACAATCTGACCAAGGCCGAGAGAGGCGGCCTGAGCGAACTGGATAAGGCCGGCTTC
ATCAAGAGACAGCTGGTGGAAACCCGGCAGATCACAAAGCACGTGGCACAGATCCTGGACTCCCGGA
TGAACACTAAGTACGACGAGAATGACAAGCTGATCCGGGAAGTGAAAGTGATCACCCTGAAGTCCAA
GCTGGTGTCCGATTTCCGGAAGGATTTCCAGTTTTACAAAGTGCGCGAGATCAACAACTACCACCACG
CCCACGACGCCTACCTGAACGCCGTCGTGGGAACCGCCCTGATCAAAAAGTACCCTAAGCTGGAAAG
CGAGTTCGTGTACGGCGACTACAAGGTGTACGACGTGCGGAAGATGATCGCCAAGAGCGAGCAGGAA
ATCGGCAAGGCTACCGCCAAGTACTTCTTCTACAGCAACATCATGAACTTTTTCAAGACCGAGATTACC
CTGGCCAACGGCGAGATCCGGAAGCGGCCTCTGATCGAGACAAACGGCGAAACCGGGGAGATCGTG
TGGGATAAGGGCCGGGATTTTGCCACCGTGCGGAAAGTGCTGAGCATGCCCCAAGTGAATATCGTGAA
AAAGACCGAGGTGCAGACAGGCGGCTTCAGCAAAGAGTCTATCCTGCCCAAGAGGAACAGCGATAA
GCTGATCGCCAGAAAGAAGGACTGGGACCCTAAGAAGTACGGCGGCTTCCTGTGGCCCACCGTGGCC
TATTCTGTGCTGGTGGTGGCCAAAGTGGAAAAGGGCAAGTCCAAGAAACTGAAGAGTGTGAAAGAG
CTGCTGGGGATCACCATCATGGAAAGAAGCAGCTTCGAGAAGAATCCCATCGACTTTCTGGAAGCCA
AGGGCTACAAAGAAGTGAAAAAGGACCTGATCATCAAGCTGCCTAAGTACTCCCTGTTCGAGCTGGA
AAACGGCCGGAAGAGAATGCTGGCCTCTGCCAAGCAGCTGCAGAAGGGAAACGAACTGGCCCTGCC
CTCCAAATATGTGAACTTCCTGTACCTGGCCAGCCACTATGAGAAGCTGAAGGGCTCCCCCGAGGATA
ATGAGCAGAAACAGCTGTTTGTGGAACAGCACAAGCACTACCTGGACGAGATCATCGAGCAGATCAG
CGAGTTCTCCAAGAGAGTGATCCTGGCCGACGCTAATCTGGACAAAGTGCTGTCCGCCTACAACAAG
CACCGGGATAAGCCCATCAGAGAGCAGGCCGAGAATATCATCCACCTGTTTACCCTGACCAATCTGGG
AGCCCCTGCCGCCTTCAAGTACTTTGACACCACCATCGACCGGAAGCAGTACAGAAGCACCAAAGAG
GTGCTGGACGCCACCCTGATCCACCAGAGCATCACCGGCCTGTACGAGACACGGATCGACCTGTCTCA
GCTGGGAGGTGACTCTGGCGGCTCAAAAAGAACCGCCGACGGCAGCGAATTCGAGCCCAAGAAGAA
GAGGAAAGTCTAA

### Supplementary Methods

**Genomic DNA extraction and SURVEYOR assay.** Genomic DNA from HEK293 cell line by transient transfection was extracted using the QuickExtract DNA Extraction Solution (Epicentre Biotechnologies). The efficiency of each individual sgRNA was tested by the SURVEYOR nuclease assay (Transgenomics) as described previously(1) using the PCR primers listed in Supplementary Table1.

### References


1. Ran FA, Hsu PD, Wright J, Agarwala V, Scott DA, and Zhang F. Genome engineering using the CRISPR-Cas9 system. *Nature Protocols.* 2013;8(11):2281-308.
